# Thermal adaptation of soil microbial growth traits in response to chronic warming

**DOI:** 10.1101/2023.05.19.541531

**Authors:** Ashley Y. Eng, Achala Narayanan, Charlotte J. Alster, Kristen M. DeAngelis

## Abstract

Adaptation of soil microbes due to warming from climate change has been observed, but it remains unknown what microbial growth traits are adaptive to warming. We studied bacterial isolates from the Harvard Forest Long-Term Ecological Research site, where field soils have been experimentally heated to 5°C above ambient temperature with unheated controls for thirty years. We hypothesized that Alphaproteobacteria from warmed plots have (1) less temperature sensitive growth rates; (2) higher optimum growth temperatures; and (3) higher maximum growth temperatures compared to isolates from control plots. We made high-throughput measurements of bacterial growth in liquid cultures over time and across temperatures from 22-37°C in 2-3°C increments. We estimated growth rates by fitting Gompertz models to the growth data. Temperature sensitivity of growth rate, optimum growth temperature, and maximum growth temperature were estimated by the Ratkowsky 1983 model and a modified Macromolecular Rate Theory (MMRT) model. To determine evidence of adaptation, we ran phylogenetic generalized least squares tests on isolates from warmed and control soils. Our results showed evidence of adaptation of higher optimum growth temperature of bacterial isolates from heated soils. However, we observed no evidence of adaptation of temperature sensitivity of growth and maximum growth temperature. Our project begins to capture the shape of the temperature response curves, but illustrates that the relationship between growth and temperature is complex and cannot be limited to a single point in the biokinetic range.

**Importance:** Soils are the largest terrestrial carbon sink and the foundation of our food, fiber, and fuel systems. Healthy soils are carbon sinks, storing more carbon than they release. This reduces the amount of carbon dioxide released to the atmosphere and buffers against climate change. Soil microbes drive biogeochemical cycling and contribute to soil health through organic matter breakdown, plant growth promotion, and nutrient distribution. In this study, we determined how soil microbial growth traits respond to long-term soil warming. We found that bacterial isolates from warmed plots showed evidence of adaptation of increased optimum growth temperature. This suggests that increased microbial biomass and growth relative to respiration in a warming world should result in greater carbon storage. As temperatures increase, greater microbial activity may help reduce the soil carbon feedback loop. Our results provide insight on how atmospheric carbon cycling and soil health may respond in a warming world.

## Introduction

The Earth’s climate is warming, and the cascading stressors from warming may have irreversible effects on microbes and the ecosystem functions that they drive. Between 2011 and 2020, earth’s land temperatures increased by 1.59°C, which is the largest rise in temperature to date (Masson-Delmotte et al., 2021). Temperature impacts rates of biological processes (Davidson & Janssens, 2006) and can result in thermal adaptation or acclimation (M. Bradford, 2013). When microbes acclimate, rates of soil processes increase less with rising temperatures compared to non-acclimated soils (M. A. Bradford et al., 2008). When microbes adapt, populations acquire new traits that fundamentally change how microbial systems respond to changes in the environment. Because soil microbes drive biogeochemical cycles and mediate atmospheric carbon fluxes (Falkowski et al., 2008; Pold et al., 2016), we need to understand the effects of long-term warming on soil microbes.

The ability of microbes to adapt to environmental change may alter ecosystem function (Wallenstein & Hall, 2012). Healthy soils are characterized by high concentrations of organic matter and abundant and active microbial communities. The activity of these microbes contributes to new organic matter deposition and soil health (Whalen et al., 2022). Soils serve as a large carbon sink, and healthy soils absorb more carbon than they release. This reduces the amount of carbon dioxide (CO_2_) emitted to the atmosphere and buffers against climate change (Stockmann et al., 2012). Changing environments may impact microbial respiration, which may also be constrained by changes in biomass and growth. Carbon storage may subsequently increase when the ratio of growth to respiration increases (Lipson, 2015). Microbial adaptation in response to warming due to climate change could thus impact soil carbon cycling (Malik et al., 2020).

To study the impacts of long-term warming on soils, a 30-year field experiment is ongoing at the Harvard Forest Long-Term Ecological Research (LTER) site in Petersham, Massachusetts. Here, experimental soils are heated 5°C above ambient temperature throughout the year since 1991 to simulate the effects of climate change. Five degrees of warming was chosen as a worst-case scenario for the rise in soil temperatures by the year 2100 (Masson-Delmotte et al., 2021); control soils received no warming treatment. Increased rates of decomposition following thirty years of warming has led to 34% loss of soil organic matter and increased flux of CO_2_ to the atmosphere in the heated versus control plots (Melillo et al., 2017). An isolate screen and metagenomic analysis showed that the ability of soil microbes to degrade complex carbohydrates also increased in response to rising temperatures (Pold et al., 2016). This was preliminary evidence of adaptation to long-term warming and suggests the potential for adaptation of other microbial traits (Melillo et al., 2017; Pold et al., 2016). Given that microbial growth and activity contribute to soil health, we sought to characterize whether microbial growth traits are associated with warming and whether they are adaptive.

We selected Alphaproteobacteria as the focus of our study because they tend to be dominant in soils, and because they showed increased absolute abundance in heated plots compared to control plots in a previous community level experiment of soil microbes (DeAngelis et al., 2015). We hypothesized that (1) Growth of Alphaproteobacteria from warmed plots are less temperature sensitive than those from control plots; (2) Optimum growth temperature of Alphaproteobacteria from warmed plots are higher than those from control plots; and (3) Maximum growth temperature of Alphaproteobacteria from warmed plots are higher than those from control plots. Given that microbes in heated soils have been exposed to higher temperatures for 22-23 years at the time of isolation, we expect them to have adapted microbial growth traits that are advantageous in warmer temperatures (Rousk et al., 2012). However, if warming does not result in adaptation of these microbial growth traits, this would suggest that changes in soil carbon dynamics may be a result of other factors such as nutrient availability, changes in microbial biomass and carbon use efficiency, or thermal acclimation (Melillo et al., 2017).

To directly measure the adaptation of bacterial growth traits due to chronic warming, we measured growth over time and across temperatures for Alphaproteobacteria isolated from the warmed and control soil plots. We estimated the intrinsic growth rate for each replicate isolate at each temperature (Zwietering et al., 1990). The Ratkowsky 1983 model (Ratkowsky et al., 1983) and a modified version of Macromolecular Rate Theory (MMRT) (Alster et al., 2022, 2023) were fitted to data for growth rate over temperature for each isolate to estimate temperature sensitivity of growth, optimum growth temperature, and maximum growth temperature. We chose the Ratkowsky 1983 model because it is a widely accepted model for bacterial growth over temperature and the MMRT model because of its underlying thermodynamic theory and application in soil microbial communities. Finally, we used phylogenetic comparative methods to test for adaptation of soil microbial growth traits (Felsenstein, 1985; Washburne et al., 2018; Yang & Rannala, 2012).

## Materials and Methods

### Isolate selection

All organisms were isolated from soils collected from the Harvard Forest long-term warming study, located at the Harvard Forest Long-Term Ecological Research (LTER) site in Petersham, MA (Peterjohn et al., 1994). The site is a mixed hardwood forest with paper and black birch (*Betula papyrifera* and lenta), red maple (*Acer rubrum*), black and red oak (*Quercus velutina* and *rubra*), and American beech (*Fagus grandifolia*) dominant tree species. Soils are coarse-loamy inceptisols. Eighteen 6 x 6 m plots were randomly assigned one of three treatments: (1) Plots with buried electrical cables, heating soils 5°C above ambient temperature throughout the year; (2) Disturbance control plots with the same as set up as the heated plots, but without electrical power; and (3) Undisturbed control plots. Soils are heated 5°C above ambient temperature by way of electrical cables buried 10 cm below the soil surface. This temperature was chosen as a worst-case scenario rise in soil temperatures by the year 2100 (IPCC, 2021).

We selected 23 strains of Alphaproteobacteria from our lab culture collection originating from either the heated or control plots. Bacteria were isolated from soils using several cultivation methods (Table S1) and cryopreserved at -80°C. Isolates were grown on 10% Tryptic Soy Agar until we could identify distinct colony morphology. We genotyped isolates by sequencing their full length 16S ribosomal RNA. We extracted genomic DNA using cetyltrimethylammonium bromide (CTAB) extraction buffer. 16S rRNA was amplified on an Eppendorf AG 22331 Hamburg using the 27F and 1492R primers. We used a 25 μl final reaction volume with 0.125 μl Invitrogen Taq, 10 μl MgCl2 10X PCR buffer, 0.75 μl 50 mM MgCl_2_, 1 μl of each primer, 2 μl of dNTP mix, and 1 μl of template for amplification reactions. We performed PCR amplifications using 35 cycles of 94°C (45 s), 50°C (30 s), 72C (120 s), followed by a final extension of 72°C (10 min). We used agarose gel electrophoresis to verify amplifications. DNA purification and Sanger sequencing was performed by Genewiz at Azenta Life Sciences.

### Genome sequencing

To extract genomic DNA for genome sequencing, strains were grown in 10% Tryptic Soy Broth, and pellets extracted using the Qiagen Blood & Tissue. We let cultures grow until late exponential phase. One day before extractions, we added 100 µl of 10% glycine to culture tubes for a final concentration of 1%; this helped to prevent cells from adhering to one another and forming clumps as cultures reached late exponential phase. DNA was eluted in TE buffer, quantified by Qubit, and transferred to the freezer for long-term storage.

Genomes were sequenced by the United States Department of Energy’s Joint Genome Institute (JGI), in house an Oxford Nanopore Technologies (ONT) MinION, or by the University of Massachusetts Medical Center (Table S1). Illumina sequencing technology was used at JGI according to standard operating procedures (Tarver et al., 2010). Long-read ONT libraries were prepared with the Ligation Sequencing Kit SQK-LSK-109 and samples were multiplexed using the Native Barcoding Expansion Kit EXP-NBD104 (Oxford Nanopore Technologies, UK). The Oxford Nanopore Native Barcoding Protocol (Oxford Nanopore Technologies, UK) was followed, and approximately 6-8 strains were multiplexed together in a run. The Covaris g-TUBE shearing step was skipped to target long fragment DNA. Starting with 1□g of DNA per strain, samples were repaired and end-prepped using the NEBNext® FFPE DNA Repair Mix and NEBNext® Ultra™ II End Repair/dA-Tailing kits (New England Biolabs, USA). DNA was cleaned using Ampure XP Beads (Beckman Coulter, USA). Samples were ligated to individual barcodes, and then about 150 ng of each sample pooled together, for a final library of 700-1000 ng. Adapters were ligated to the sample with Blunt/TA ligase (New England Biolabs, USA). The long fragment buffer provided in the sequencing kit was used in an extended 10 minute incubation at 37℃ to enrich for high-molecular weight DNA. The flow cell was primed using the Flow Cell Priming Kit (Oxford Nanopore Technologies, UK), and about 15 fmols of the library was mixed with the sequencing buffer and loading beads and then loaded through the Spot-On port of the flow cell (Choudoir et al., 2023).

Sequence runs were initially basecalled using the high-accuracy base calling (HAC) algorithm with Guppy (R. R. Wick et al., 2019). These fast5 files were concatenated into one file, then reads were subsampled based on read quality using Filtlong (R. Wick & Menzel, 2019). The filtered reads were *de novo* assembled using Flye (Kolmogorov et al., 2019). A consensus assembly was generated using Racon (Vaser et al., 2017), and final polishing performed with Medaka (*Medaka*, 2017/2023). These assemblies were checked for quality using Quast (Gurevich et al., 2013) and CheckM (Parks et al., 2015).

### Quantifying microbial growth in liquid culture

We measured bacterial growth over time using absorbance measurements for liquid cultures spanning temperatures from 22-37°C along 2-3°C increments. We chose 22-37°C to capture the temperature growth range of mesophiles and to accommodate instrumental limitations. Absorbance was measured in 96-well plates by optical density at 600 nm (OD_600_ nm) as a measure of cell abundance. Each well was filled with 240 μl of 10% Tryptic Soy Broth. Colonies grown on petri plates were resuspended in 500 μl of Phosphate Buffered Saline and 10 μl resuspension was inoculated into each well. For temperatures at or above 30°C, we pipetted 2-3 ml of a 0.05% solution of Triton X-100 in 20% ethanol on the plate lid to prevent condensation. Each plate accommodated 8 isolates with 11 replicates each and 8 negative controls according to a randomized plate format. Bacterial growth was measured using a SpectraMax M2 plate reader (Molecular Devices, CA) at OD_600_ nm. Growth curves lasted 72-99 hours or until microbes entered death phase.

### Model fitting

We fitted the Gompertz growth curve on data for OD_600_ nm over time to calculate growth rate using the R package Growthcurver (Sprouffske & Wagner, 2016) (figure 1A). A Gompertz growth curve is an established time course model that parametrizes bacterial growth over time as a sigmoidal function (Zwietering et al., 1990). We estimated the intrinsic growth rate from the fitted Gompertz model. This was repeated for each replicate at each temperature.

**Fig 1.**
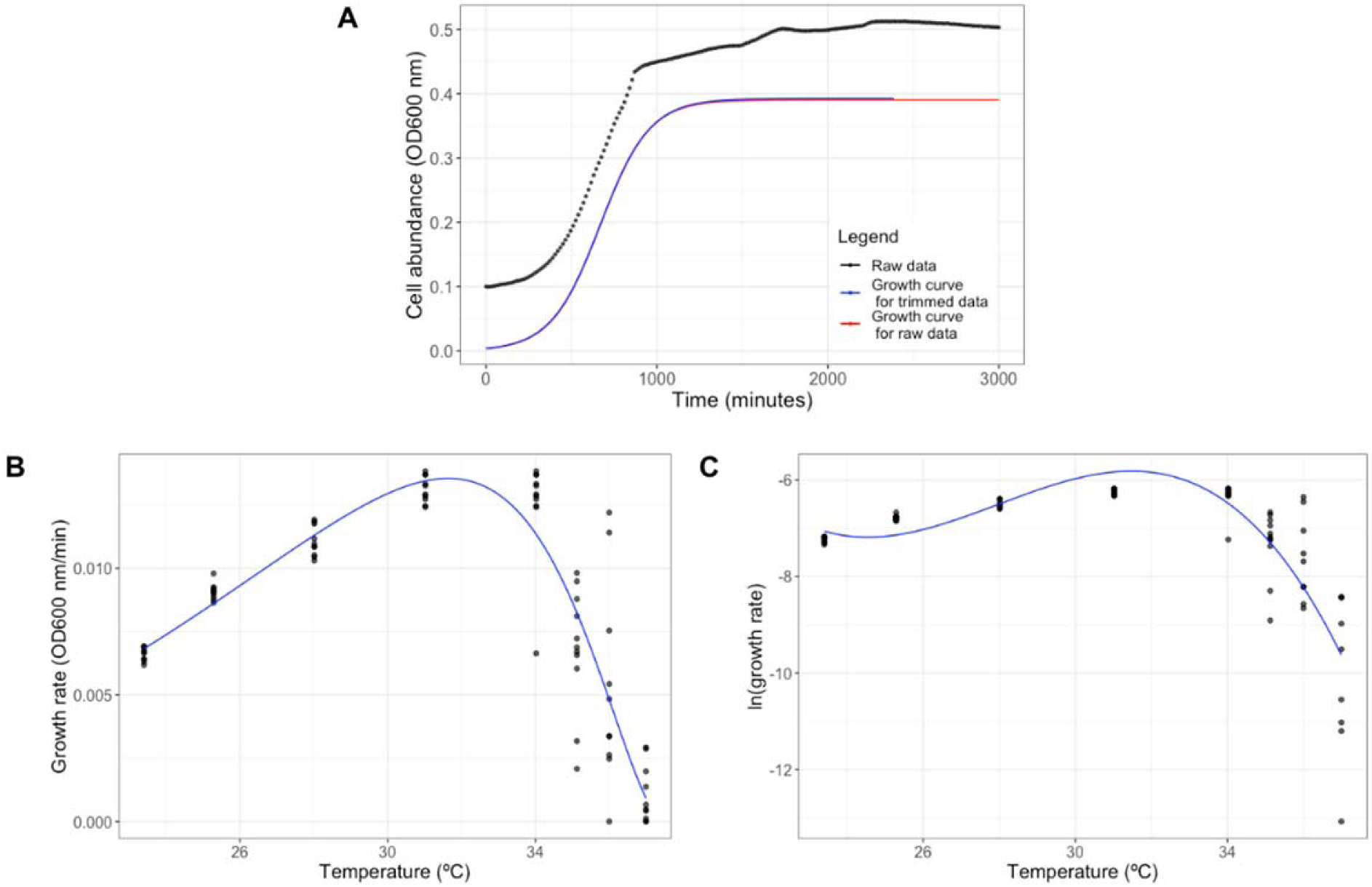
Example data sets for measuring growth parameters and estimating temperature sensitivity of growth using two different models. (A) Cell abundance over time was measured for liquid growth curves between 22-37°C in 2-3°C increments. A Gompertz growth curve was fit on data of cell abundance over time. Intrinsic growth rate was extracted from the fitted model. (B) The Ratkowsky 1983 model was fitted to growth rate over temperature for each isolate. Temperature sensitivity of growth, optimum growth temperature, and maximum growth temperature were estimated from the fitted model. (C) A modified version Macromolecular Rate Theory was fitted to natural log-transformed data of growth rate over temperature for each isolate. Values of zero were removed from analysis. Temperature inflection point and optimum growth temperature were calculated from the fitted model. Data for A, B, and C are of the same isolate.

We fitted temperature response curves to estimate the microbial growth traits.

Temperature sensitivity of growth is an estimate of how growth rate changes with increasing temperature, but it can be estimated using different parameters depending on the model applied (Table 1). Optimum growth temperature is the temperature at which growth rate is the greatest. Maximum growth temperature is the estimated highest temperature at which microbial growth is permissible. We modeled the relationship between growth rate and temperature for each isolate using the Ratkowsky 1983 model (figure 1B):

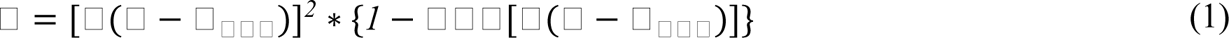

**Table 1.**
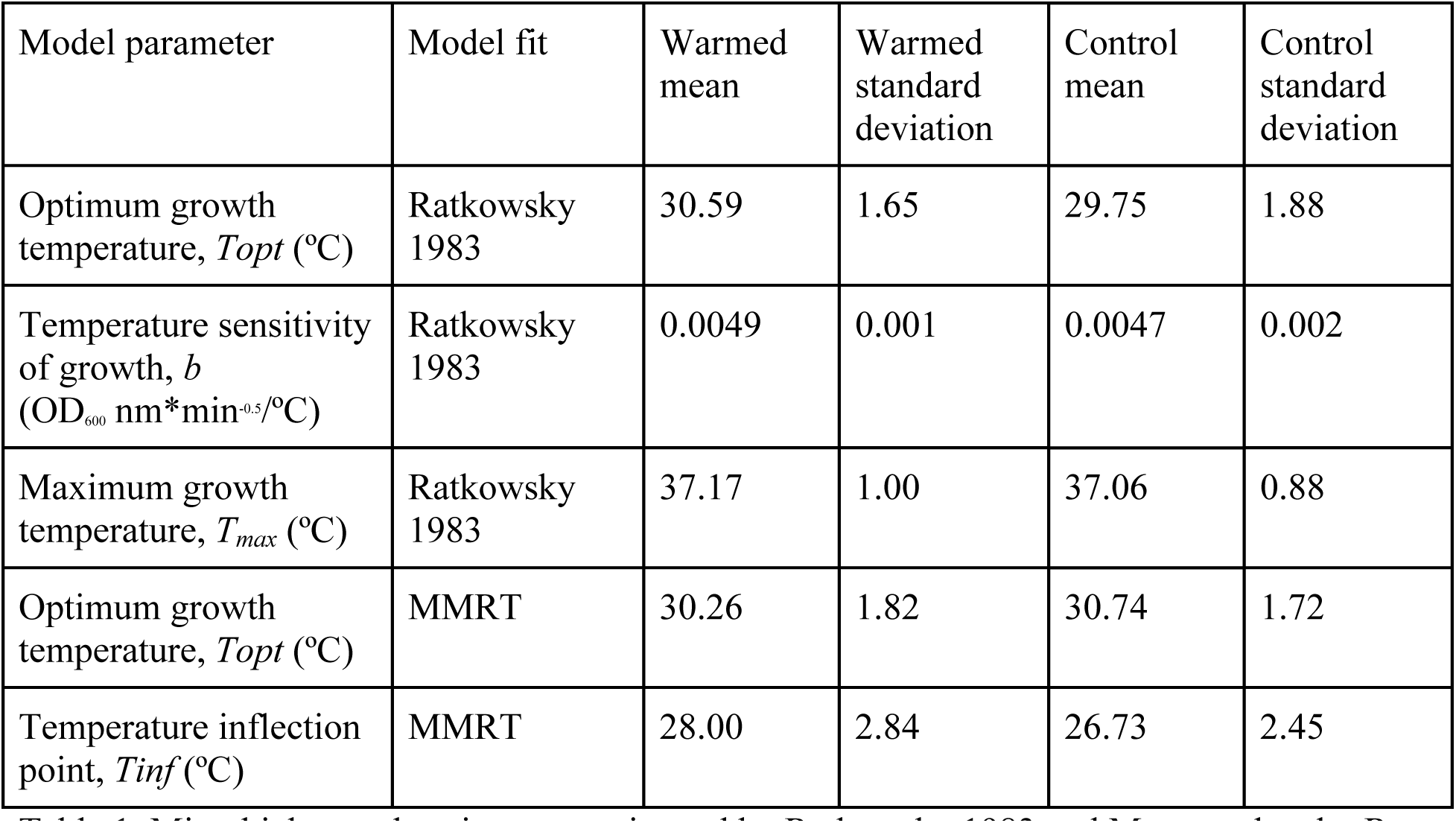
Microbial growth traits were estimated by Ratkowsky 1983 and Macromolecular Rate Theory (MMRT) model parameters. The mean and standard deviation of model parameters for isolates from the warmed and control plots were also calculated.

| Model parameter | Model fit | Warmed mean | Warmed standard deviation | Control mean | Control standard deviation |
| --- | --- | --- | --- | --- | --- |
| Optimum growth temperature, $T_{opt}$ (°C) | Ratkowsky 1983 | 30.59 | 1.65 | 29.75 | 1.88 |
| Temperature sensitivity of growth, $b$ ($OD_{600} \text{ nm} \cdot \text{min}^{-0.5} / ^\circ\text{C}$) | Ratkowsky 1983 | 0.0049 | 0.001 | 0.0047 | 0.002 |
| Maximum growth temperature, $T_{max}$ (°C) | Ratkowsky 1983 | 37.17 | 1.00 | 37.06 | 0.88 |
| Optimum growth temperature, $T_{opt}$ (°C) | MMRT | 30.26 | 1.82 | 30.74 | 1.72 |
| Temperature inflection point, $T_{inf}$ (°C) | MMRT | 28.00 | 2.84 | 26.73 | 2.45 |

*T_min_* is the minimum permissible temperature for growth (°C), *T_max_* is the maximum permissible growth temperature (°C), and *c* is an empirical parameter required to model data above the optimum temperature (°C^-1^). Temperature sensitivity of growth was quantified by the Ratkowsky parameter *b* (OD_600_nm*min^-0.5^/°C), which is the regression coefficient for square root of growth rate on temperature (Ratkowsky et al., 1983; Zwietering et al., 1991). Maximum growth *T_max_*(°C) was extracted as a parameter from the fitted model. Optimum growth *Topt* (°C) was estimated from the fitted model (Padfield et al., 2021). Model fitting was performed using the R package *rTPC* (Padfield et al., 2021).

The temperature optima (*Topt*) and inflection point (*Tinf*) of each bacterial growth curve were estimated using a modified version of Macromolecular Rate Theory (MMRT) (figure 1C):

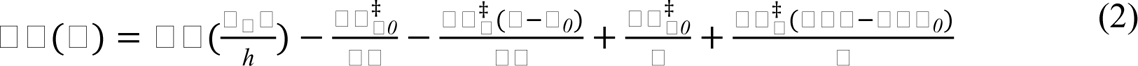

where, *k* is the bacterial growth rate (OD_600_nm), *k*_B_ is Boltzmann’s constant, T is temperature (K), *h* is Planck’s constant, R is the universal gas constant, 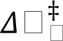 (superscript denotes transition state) is the change in enthalpy (J mol^-1^), 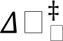 is the change in entropy (J mol^-1^ K^-1^), 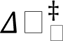 is the change in heat capacity (J mol^-1^ K^-1^), and T_0_ is the reference temperature (set to 296.1K) (Hobbs et al., 2013). The MMRT equation was modified to allow 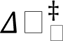 to vary linearly with temperature:

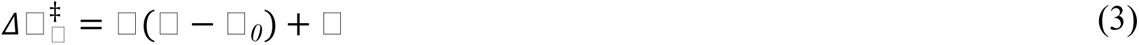

where, *A* is the slope and *B* is the value of 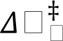 at *T_0_* (Alster et al., 2023). We used a non-linear least-squares regression in R version 4.2.1 (R Core Team 2021) to fit MMRT and calculated the *Topt* and *Tinf* numerically using the first and second derivative, respectively. We chose to use the modified version of MMRT to better capture the *Topt* due to asymmetries observed in the temperature response data and because 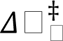 varies over a wide temperature range (Darros-Barbosa et al., 2003; Ghosh & McSween Jr., 1999; Prentice et al., 2020). We calculated residual standard errors to determine whether the modified MMRT or original MMRT more adequately fit our data (Table S2).

To evaluate the fit of the models, we calculated the residual standard error (RSE). The Ratkowsky parameter *(b)*, *Topt* estimated by both Ratkowsky 1983 and MMRT, *Tmax* estimated by Ratkowsky 1983, and *Tinf* estimated by MMRT were used as traits for the phylogenetic group comparison. Outliers, potentially due to irregularities in the replicate, were excluded from analysis if they were not within the same order of magnitude as the remaining points in the data set.

### Phylogenetic group comparison

We conducted a phylogenetic group comparison of traits (Felsenstein, 1985; Washburne et al., 2018; Yang & Rannala, 2012) test our hypotheses that Alphaproteobacteria from warmed plots have (1) less temperature sensitive growth rates; (2) higher optimum growth temperatures; and (3) higher maximum growth temperatures compared to isolates from control plots. We used the R package *nlme* to conduct phylogenetic generalized least squares (PGLS) test (Garland Jr. et al., 1993; Lindstrom & Bates, 1990). A phylogenetic group comparison accounts for the lack of independence in phylogenetic hierarchical species data. A Lilliefors test for normality was used to determine whether residuals of PGLS tests were normally distributed, and Q-Q plots were made. Trait data was transformed if the Lilliefors test failed. We removed outlier data points, which were 3-4 orders of magnitude greater than the remaining points in the data set.

A genome-based phylogeny was constructed using the United States Department of Energy’s Systems Biology Knowledgebase (KBase) (Arkin et al., 2018). We quality checked genomes using the Quast (Gurevich et al., 2013) and CheckM (Parks et al., 2015) applications in KBase. Genomes were annotated by Prokka (Seemann, 2014). The phylogeny was constructed using Insert Genome into SpeciesTree v2.2.0 (Price et al., 2009), which creates a multiple sequence alignment based on universal genes defined by COG (Clusters of Orthologous Groups) gene families. We set the nearest public genome count to one and removed the public node in R using the package *ape* (Paradis & Schliep, 2019). Temperature sensitivity of growth, optimum growth temperature, and maximum growth temperature were mapped as traits on the phylogeny.

We also calculated phylogenetic signal using Pagel’s λ to quantify the tendency of the isolates to closely resemble each other based on their phylogenetic distribution of traits. Pagel’s λ is a measure of correlation between species under Brownian Motion (Pagel, 1999) and was estimated using the R package *phytools* (Revell, 2012). P-values and group means were calculated for each microbial growth trait from warmed and control soils. Soil microbial isolates were the experimental unit, and all P-values less than 0.05 were considered statistically significant relationships. All statistical analyses were performed in R using RStudio (*RStudio Team,* version 20200.02.0+443).

## Results

### Optimum growth temperature quantified by Ratkowsky 1983 model shows evidence of adaptation

There was a significant difference between optimum growth temperature (*Topt*) of isolates from heated versus control plots quantified by the Ratkowsky 1983 model (*t*(23) = 2.84, *p* = 0.01, n_warm_ = 8, n_control_ = 15) (Table S3). Optimum growth temperature for isolates from warmed plots (*M* = 30.59, *SD* = 1.65) was greater than those of control plots (*M =* 29.75, *SD* = 1.88). Residuals were normally distributed according to the Lilliefors test (*p* = 0.41, n_warm_ = 8, n_control_ = 15). Pagel’s λ showed that *Topt* was not distributed according to Brownian motion (λ = 6.61E-05, *p* = 1.00) (figure 2A).

**Fig 2.**
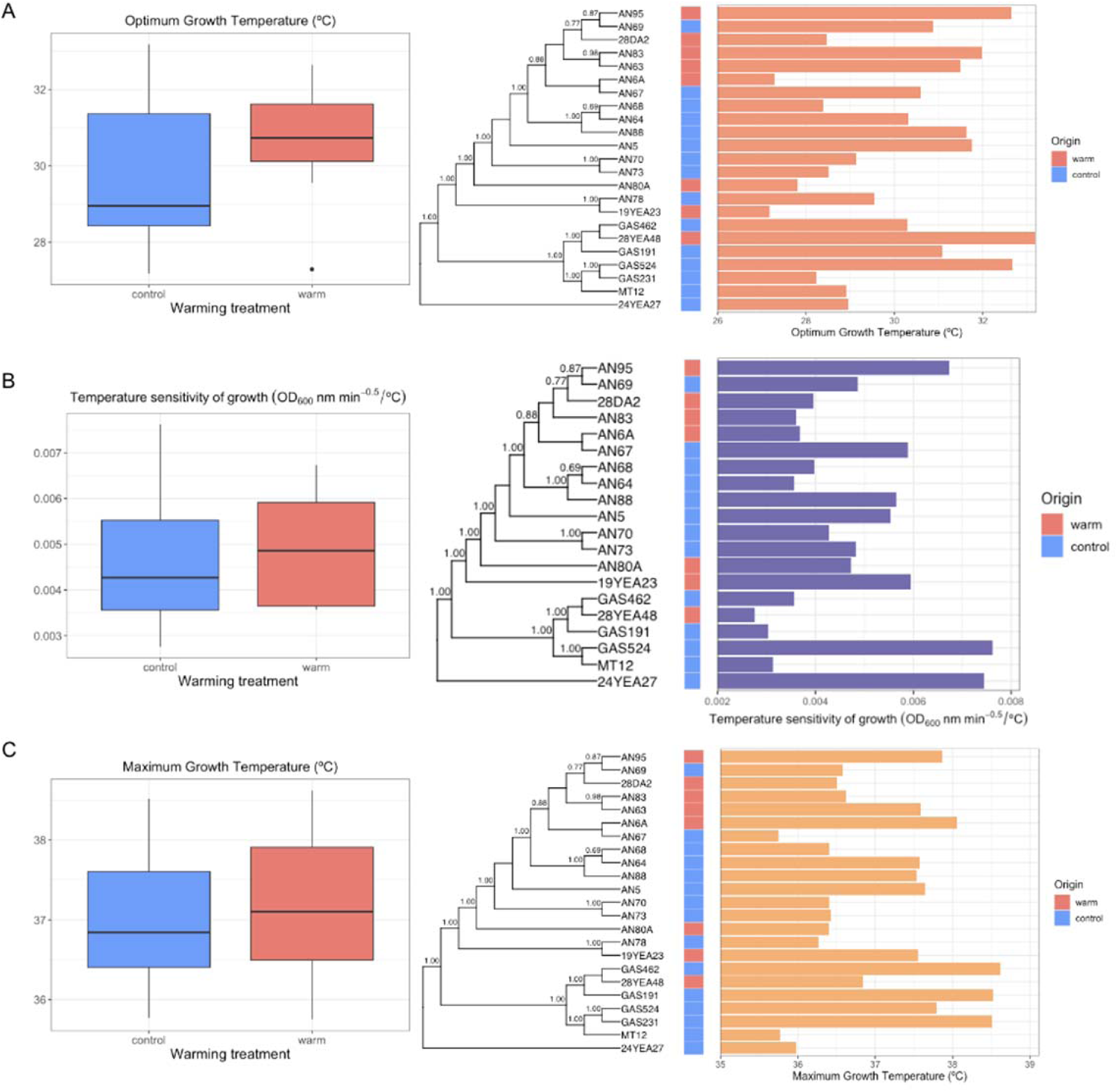
Optimum growth temperature (A), temperature sensitivity of growth (B), and maximum growth temperature (B) were quantified by fitting the Ratkowsky 1983 model on data for growth rate over temperature for each isolate. A multiple sequence alignment of universal genes found in core orthologous group genes was used to construct the phylogenetic tree. Phylogenetic generalized least squares test was used to test for difference in optimum growth temperature between isolates from heated and control soil plots.

### No evidence of adaptation of temperature sensitivity of growth quantified by Ratkowsky 1983

Our results showed no significant difference in temperature sensitivity of growth quantified by the Ratkowsky parameter (*t*(20) = 1.79, *p* = 0.09, n_warm_ = 7, n_control_ = 13) (Table S3). Temperature sensitivity of isolates from warmed and control plots were 0.0049 (*SD* = 0.001) and 0.0047 (*SD* = 0.002) respectively. Residuals were normally distributed according to the Lilliefors test (*p >* 0.05). Pagel’s λ showed that temperature sensitivity of growth was not distributed according to Brownian motion, suggesting a random distribution (λ = 6.61E-05, *p* = 1.00) (figure 2B).

### No evidence of adaptation of maximum growth temperature quantified by Ratkowsky 1983

Results of the phylogenetic least squares test for maximum growth temperature quantified by the Ratkowsky 1983 model showed no significant difference between isolates from heated and control plots (*t*(23) = -0.35, *p* > 0.05, n_warm_ = 8, n_control_ = 15) (Table S3). Maximum growth temperature for isolates from the warmed and control plots were 37.17°C (*SD* = 1.00) and 37.06°C (*SD* = 0.88) respectively. Residuals were normally distributed according to the Lilliefors test (*p* = 0.45). Pagel’s λ showed that maximum growth temperature was not distributed according to Brownian motion (λ = 6.61E-05, *p* = 1.00) (figure 2C).

### Temperature sensitivity inferred by Macromolecular Rate Theory does not show evidence of adaptation

Results of the phylogenetic generalized least squares test for optimum growth temperature quantified by MMRT showed no significant difference between isolates from the heated and control plots (*t*(23) = 1.60, *p* = 0.12, n_warm_ = 8, n_control_ = 15) (Table S4). Optimum growth temperature for isolates from warmed plots (*M* = 30.26, *SD =* 1.82) was not significantly different than those of control plots (*M* = 30.74, *SD =* 1.72). Residuals were normally distributed according to the Lilliefors test (*p* > 0.05). Pagel’s λ showed that *Topt* was not distributed according to Brownian motion (λ = 0.45, *p* > 0.05) (figure 3). Residual standard errors indicated that the Ratkowsky 1983 model more adequately fit our data in comparison to the MMRT model (Table S2).

**Fig 3.**
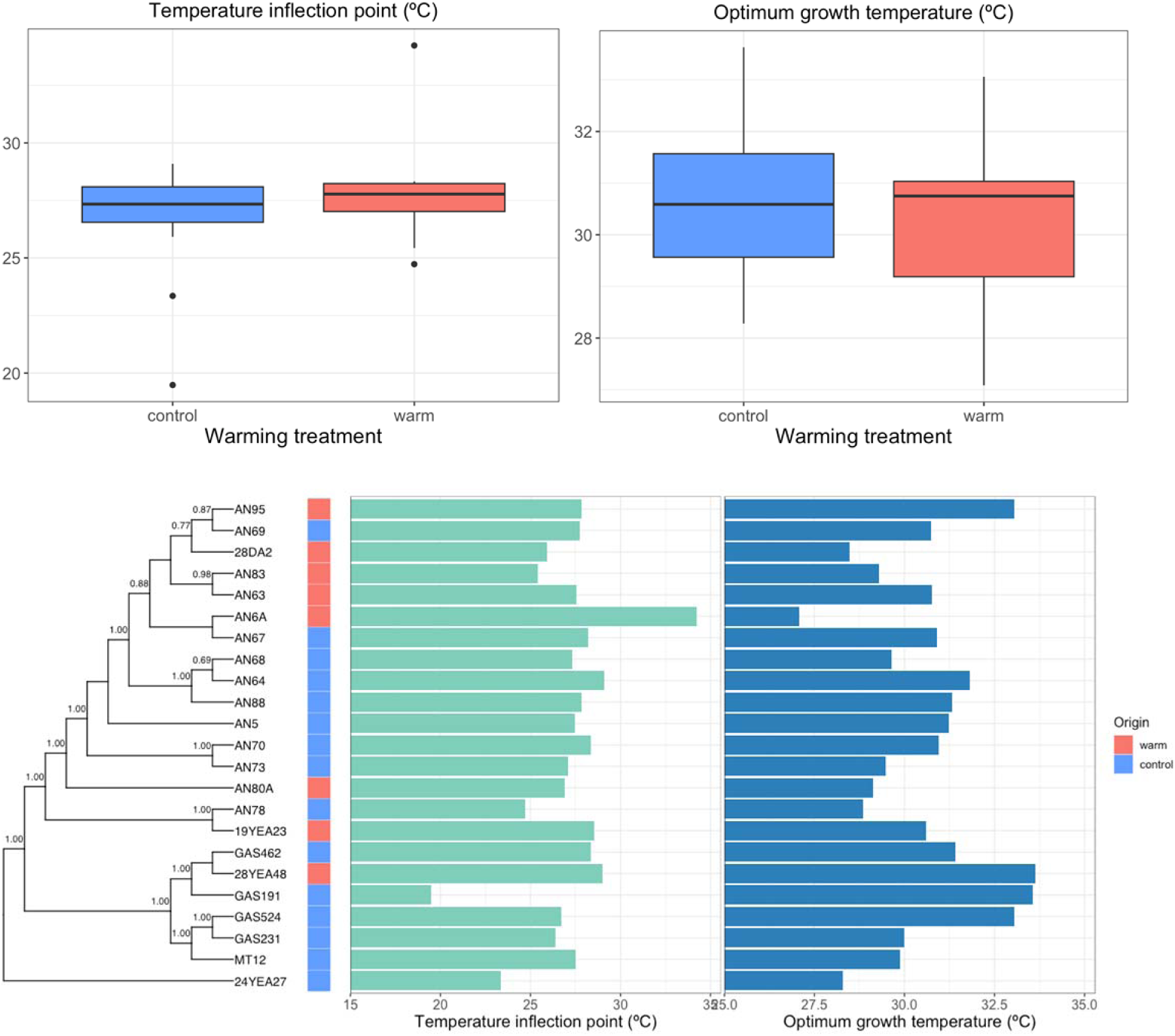
Temperature inflection point (*Tinf*) and optimum growth temperature (*Topt*) were quantified through fitting the Macromolecular Rate Theory model on data for growth rate over temperature for each isolate. A multiple sequence alignment of universal genes found in core orthologous group genes was used to construct the phylogenetic tree. Phylogenetic generalized least squares test was used to test for difference in *Tinf* and *Topt* between isolates from heated and control soil plots.

There was no significant difference in temperature inflection point (*Tinf*) between isolates from the heated and control soils (*t*(23) = 1.04, *p* > 0.05, n_warm_ = 8, n_control_ = 15) (Table S4). Residuals of PGLS on untransformed *Tinf* data failed the Lilliefors test for normality (P < 0.05). Average temperature inflection point for isolates from warmed plots was 28.00°C (*SD =* 2.84). Average temperature inflection point for isolates from control plots was 26.73°C (*SD =* 2.45). Residuals failed the Lilliefors test for normality following log, square root, and lambda (Box-Cox test) transformations (*p* < 0.05). Pagel’s λ showed that *Tinf* was not distributed according to Brownian motion (λ = 6.61E-05, *p* > 0.05) (Figure 3). Since the residuals failed the Lilliefors test for normality, we transformed *Tinf* (i.e. log transformation, square root, box-cox). However, all transformations also failed the test for normality, and suggest that results of PGLS for *Tinf* estimated by MMRT should be interpreted with caution.

## Discussion

We expected Alphaproteobacteria isolated from warmed plots would have (1) lower temperature sensitivities of growth; (2) higher optimum growth temperatures; and (3) higher maximum growth temperatures compared to isolates from control plots. Our results showed evidence of adaptation of optimum growth temperature quantified by the Ratkowsky 1983 model, but not for other measured traits. Evidence of adaptation of *Topt* estimated by the Ratkowsky 1983 model affirm observations from previous studies, where increased optimum growth temperature is associated with warmer soils (Donhauser et al., 2020; Smith et al., 2022). However, the lack of differences observed in other microbial growth traits estimated by both the Ratkowsky 1983 and MMRT models may be due to the shape of the temperature response curve, model fitting, or the magnitude and duration of warming, for example. Evidence for this conclusion lies in the observation that the Ratkowsky 1983 model fit was better than the MMRT model for optimum growth temperature.

The difference in evidence of adaptation for optimum growth temperature quantified by the Ratkowsky 1983 and modified MMRT models may be due to a difference in fits. The residual standard errors for the Ratkowsky 1983 fitted model on each isolate is 2-3 orders of magnitude lower than those of the MMRT fitted models (Table S2). Adequate fitting of MMRT requires a dataset to at least capture the optimum growth temperature. Although our dataset includes *Topt*, it is considerably limited at lower temperatures and lacks growth rate data at the temperature minima. This limitation may be associated with the less accurate MMRT fits, resulting in inaccurate estimations of *Topt*. The difference in evidence of adaptation when *Topt* was estimated by the Ratkowsky 1983 and MMRT models demonstrate the importance of model fit when estimating microbial traits.

There are several differences between the Ratkowsky 1983 model and Macromolecular Rate Theory. Ratkowsky 1983 is an empirically determined model of growth rate over temperature for each isolate (Ratkowsky et al., 1983). MMRT is based on thermodynamic theory and is not empirically determined. It accounts for changes in the temperature response in the absence of enzyme denaturation at temperatures above the optimum temperature through changes in heat capacity. The residual standard errors indicate that the Ratkowsky 1983 is a more appropriate fit for our data of growth rate over temperature compared to MMRT. However, we are particularly interested in MMRT due to its underlying thermodynamic theory, as well as its application in soil ecosystems (Alster et al., 2020, 2022, 2023).

The lack of evidence of adaptation of other microbial growth traits demonstrates the limitations of inferring microbial growth traits based on a single temperature point. Traits such as temperature sensitivity of growth are more nuanced and may be impacted by thermal niche breadth. Thermal niche breadth is the range of temperatures that permits microbial growth. Previous studies observed that changes in the range between minimum and maximum growth temperatures depended on soil incubation temperatures (Rijkers et al., 2022; van Gestel et al., 2013). This suggests that the relationship between growth rate and temperature may also vary between minimum and maximum growth temperatures. The rate at which growth rate changes across temperatures, or the steepness of the temperature response curve, may be impacted by the environment, thus altering thermal niche breadth. Challenges in quantifying change in microbial growth rate over temperature may result if such environmental factors are not fully accounted for. This concept of a thermal niche breadth may have an associated fitness cost, as seen with other microorganisms (Herren & Baym, 2022). Therefore, it may be challenging to identify microbial growth trait adaptation without also considering changing thermal niche breadths.

Pagel’s λ was intermediate (0 < λ < 1) for all microbial growth traits, which suggests that the distribution of traits was not as expected under Brownian Motion. There are multiple explanations for such results. One explanation is that climate warming may be associated with selection of intermediate phenotypes (i.e. stabilizing selection) instead of extremes. This may have resulted in constrained trait evolution. Additionally, changes in evolutionary rate over time may have also resulted in non-Brownian Motion distribution of traits (Smith et al., 2022a). It is possible that discontinuous substrate availability over the decades of experimental warming could have caused a difference in growth rate, and possibly evolutionary rate, over time (Melillo et al., 2017). Phylogenetic signal is also often quantified by Blomberg’s *K*, which is a variance ratio and has the advantage of being able to be greater than one. However, our data was not suitable for Blomberg’s *K* estimations as it resulted in a singular matrix.

Soil microbial growth tends to be limited by substrate availability, so evidence of adaptation from PGLS tests may have been occluded by high levels of nutrient availability in the laboratory growth conditions of these experiments. Kamble *et al*. (2018) observed that bacterial and fungal growth in soils was carbon limited. In a community level experiment in a boreal forest, Ekblad *et al*. (2002) observed that soil microbial biomass was limited by carbon but not nitrogen availability. Although these experiments were conducted in soils on the community level, it is possible that the carbon and nutrient rich media used in this study may obscure the effect of nutrient availability and substrate-specific growth dynamics of microbes in warming soils. Studying microbial growth under lower nutrient conditions may provide a different perspective on how warming impacts microbial growth traits.

Thermal adaptation of increasing growth with temperature has been observed for other organisms in response to climate warming. Among other microorganisms, growth rate of pathogenic fungi, *Mycosphaerella graminicola* was observed to be associated with increasing temperatures (Zhan & McDonald, 2011). Globally distributed plant pathogens were also found to locally adapt to their environments, resulting in significantly different optimum growth temperatures (Boixel et al., 2022). Thermal adaptation is also often investigated more broadly among other ectotherms. Villeneuve *et al*. (2021) observed that growth of *Urosalpinx cinerea* (Atlantic oyster drill) was positively associated with spawning temperature. Studying thermal adaptation is highly relevant as the effects of the climate crisis increase. However, doing so is challenging among organisms with longer generation times, which highlights the importance of utilizing techniques beyond lab and field-based experiments and suggests a benefit to studying adaptation among organisms with short generation times and large populations like microbes. It is also possible that the organismic adaptation to temperature appears overly significant due to the difficulty in publishing negative or non-significant results.

Change in microbial growth traits is just one example of how warming may impact soil microbes. Increasing temperatures are also associated with evolutionary selection of organisms with smaller genome sizes, as seen in fire-affected soils (Sorensen et al., 2019). Evidence of adaptation for other metabolic processes, such as respiration, has also been observed (Nottingham et al., 2022; Tian et al., 2022). Differences in microbial growth traits between isolates from warmed and control soils may be due to reasons other than adaptation. Such differences may be due to depletion of labile carbon (M. A. Bradford et al., 2008; Melillo et al., 2017), changing microbial community structure (Frey et al., 2008; Melillo et al., 2017), microbial physiology (Allison et al., 2010), and species sorting and functional diversity (Smith et al., 2022).

While thermal adaptation of microbial traits has been observed in other studies, our results demonstrate that measuring growth potential may be impacted by additional factors. We used laboratory settings to quantify microbial growth traits, which may be an inaccurate representation of field conditions. Under these conditions, results of our study suggest that warming has not resulted in adaptation of temperature sensitivity of growth and maximum growth temperature quantified by the Ratkowsky 1983 model and temperature inflection point and optimum growth temperature quantified by MMRT. However, optimum growth temperature estimated by Ratkowsky 1983 showed some evidence of adaptation. As temperatures increase, changes in soil microbial growth rate may affect rates of atmospheric carbon cycling. Future exploration of whether growth strategies explain microbial adaptation to warming will help predict changes in microbial community and ecosystem function and allow us to better understand soil microbial responses to warming.

## Data availability

The whole genome assemblies have been deposited at GenBank under the accession numbers listed in Table S1, along with the raw data deposited in the Sequence Read Archive, and associated BioProject and BioSamples. Microbial growth data can be found in the Harvard Forest Data archive [HF data accession number]. All code used for model fitting and data analysis are available in the supplemental information.

## Acknowledgements

This project was supported by a grant from the National Science Foundation (No. DEB-1749206) to KMD. The soil warming experiments at Harvard Forest are maintained with support from the National Science Foundation (NSF) Long Term Ecological Research Program (DEB-1832110) and a Long-Term Research in Environmental Biology grant (DEB-1456610). We are grateful to Serita Frey and Mel Knorr for their support of our soil sampling. The work conducted by the U.S. Department of Energy Joint Genome Institute (https://ror.org/04xm1d337), a DOE Office of Science User Facility, is supported by the Office of Science of the U.S. Department of Energy operated under Contract No. DE-AC02-05CH11231. We are grateful to all the people who contributed to the isolation, genotyping, sequencing, and annotation of isolates in this study, including (but probably not limited to) Erin Bergeron, Andrew Billings, Isabella Bushko, Gina Chaput, Mallory Choudoir, Emily Clark, Luiz Domeignoz-Horta, Alon Efroni, Spencer Moore, Samantha Murphy, Grace Pold, Damayanti Rodriguez-Ramos, Rachel Simoes, Abigail Sondrini, Bianca Surjawan, and Wing Yin Tam.

